# Improving preterm newborn identification in low-resource settings with machine learning

**DOI:** 10.1101/334904

**Authors:** Katelyn J. Rittenhouse, Bellington Vwalika, Alex Keil, Jennifer Winston, Marie Stoner, Joan T. Price, Monica Kapasa, Mulaya Mubambe, Vanilla Banda, Whyson Muunga, Jeffrey S.A. Stringer

## Abstract

**Background:** Globally, preterm birth is the leading cause of neonatal death with estimated prevalence and associated mortality highest in low‐ and middle‐income countries (LMICs). Accurate identification of preterm infants is important at the individual level for appropriate clinical intervention as well as at the population level for informed policy decisions and resource allocation. As early prenatal ultrasound is commonly not available in these settings, gestational age (GA) is often estimated using newborn assessment at birth. This approach assumes last menstrual period to be unreliable and birthweight to be unable to distinguish preterm infants from those that are small for gestational age (SGA). We sought to leverage machine learning algorithms incorporating maternal factors associated with SGA to improve accuracy of preterm newborn identification in LMIC settings.

**Methods and Findings:** This study uses data from an ongoing obstetrical cohort in Lusaka, Zambia that uses early pregnancy ultrasound to estimate GA. Our intent was to identify the best set of parameters commonly available at delivery to correctly categorize births as either preterm (<37 weeks) or term, compared to GA assigned by early ultrasound as the gold standard. Trained midwives conducted a newborn assessment (<72 hours) and collected maternal and neonatal data at the time of delivery or shortly thereafter. New Ballard Score (NBS), last menstrual period (LMP), and birth weight were used individually to assign GA at delivery and categorize each birth as either preterm or term. Additionally, machine learning techniques incorporated combinations of these measures with several maternal and newborn characteristics associated with prematurity and SGA to develop GA at delivery and preterm birth prediction models. The distribution and accuracy of all models were compared to early ultrasound dating. Within our live‐born cohort to date (n = 862), the median GA at delivery by early ultrasound was 39.4 weeks (IQR: 38.3 ‐ 40.3). Among assessed newborns with complete data included in this analysis (n = 458), the median GA by ultrasound was 39.6 weeks (IQR: 38.4 ‐ 40.3). Using machine learning, we identified a combination of six accessible parameters (LMP, birth weight, twin delivery, maternal height, hypertension in labor, and HIV serostatus) that can be used by machine learning to outperform current GA prediction methods. For preterm birth prediction, this combination of covariates correctly classified >94% of newborns and achieved an area under the curve (AUC) of 0.9796.

**Conclusions:** We identified a parsimonious list of variables that can be used by machine learning approaches to improve accuracy of preterm newborn identification. Our best performing model included LMP, birth weight, twin delivery, HIV serostatus, and maternal factors associated with SGA. These variables are all easily collected at delivery, reducing the skill and time required by the frontline health worker to assess GA.

## INTRODUCTION

Preterm birth affects more than one in ten live births worldwide.[1] It is the single largest cause of neonatal death and the second leading cause of death in children under the age of 5 years.[2] Many babies who survive a preterm birth face life‐long morbidity, including cognitive disability, poor motors skills, behavioral problems, hearing loss, chronic lung disease, and decreased economic productivity.[3–5] The greatest burden of preterm birth falls on low‐and middle‐income countries (LMICs), where more than 90% of the global 15 million preterm deliveries occur each year[6] and where preterm infants carry a 7‐fold higher risk of neonatal mortality and a 2.5‐fold higher risk of post‐neonatal mortality compared to their full‐term counterparts.[7] In these settings, preterm infants often go unrecognized due to inaccurate estimation of gestational age (GA). On the individual level, this can result in missed opportunities for clinical intervention; on the population level, this can limit the ability to monitor preterm birth rates and make informed decisions around policy and resource allocation.

Early prenatal ultrasound, widely regarded as the gold standard for GA dating, is unavailable in many LMIC settings. In its absence, providers must rely on other methods, such as last menstrual period (LMP), newborn assessment, or birthweight to classify infant GA at delivery. Each of these approaches has limitations. Reported LMP is subject to patient recall and can be very unreliable in settings where women present late for care.[8–11] Newborn assessment, including the commonly used New Ballard Score (NBS),[12] suffers from poor inter‐rater reliability[13, 14] and tends to overestimate GA, particularly in LMICs[15] and settings with high rates of small‐for‐gestational age (SGA).[16–21] Finally, birthweight, while an easily obtained and reliable indicator, does not distinguish between an infant that is preterm and one that is SGA.

We sought to develop a machine learning algorithm that can estimate GA at birth from readily obtained indicators in a setting where early ultrasound is not available. We were particularly interested in the simple, binary classification of preterm (i.e., <37 weeks) versus term. We hypothesized that a model combining LMP, individual elements of the NBS, birthweight, and key pregnancy risk factors associated with SGA, would outperform any individual approach.

## METHODS

This study was conducted using data from the Zambian Preterm Birth Prevention Study (ZAPPS; ClinicalTrials.gov identifier: NCT02738892), an ongoing prospective obstetrical cohort at the Women and Newborn Hospital of the University Teaching Hospital (UTH) in Lusaka, Zambia. The rationale for our study, its procedures, and cohort characteristics have been described elsewhere.[22] Briefly, women are enrolled in early pregnancy and followed through delivery and the postpartum period. Written informed consent is obtained from all participants prior to study enrollment for collection of maternal and newborn data. GA is established by ultrasound (Sonosite M‐Turbo; Fuji Sonosite, Inc, Bothell, WA) at study screening using the fetal crown rump length (if <14 weeks gestation)[23] or head circumference and femur length (if ≥14 weeks).[24] All fetal biometry measurements are measured twice and then averaged.

The study employs midwives who attend to participants admitted to the labor ward or postpartum unit at UTH. Their duties include ensuring that relevant clinical information is captured in the study record, that babies are weighed at birth or shortly thereafter, and that the NBS is performed within 72 hours of delivery.

Newborns were included in this analysis if they were live‐born and they had a complete set of characteristics and metrics assessed in this study. We defined preterm birth as birth prior to 37 weeks of gestation and SGA as a birthweight less than the 10^th^ percentile for its corresponding GA.[25] The NBS sums assessments of 5 domains of neuromuscular maturity and 7 domains of physical maturity into a composite score that is used to assign GA at delivery.[12] We evaluated both the composite score and its individual components in this study.

In our analyses, we assessed eight models: three single parameter GA dating methods and five multiple parameter novel machine learning GA dating models (Table 1). We were primarily interested in classifying preterm birth as a binary outcome (i.e., <37 weeks or not), but we also wished to assess how the models might estimate GA as a continuous outcome. We restricted our models to maternal and newborn characteristics that are accessible to health workers in resource‐limited settings at the time of delivery, either through direct assessment or review of the medical record.

**Table 1:** Components of gestational age dating models

| GA Dating Model | GA at delivery (composite NBS) | GA at delivery (LMP) | GA at delivery (birth weight 50%ile)<sup>^</sup> | NBS (individual components)<sup>†</sup> | Birth weight | Maternal height | Hypertension in labor | Maternal HIV infection | Twin gestation |
| --- | --- | --- | --- | --- | --- | --- | --- | --- | --- |
| NBS | • |  |  |  |  |  |  |  |  |
| LMP |  | • |  |  |  |  |  |  |  |
| Birth weight |  |  | • |  |  |  |  |  |  |
| Optimized NBS |  |  |  | • |  |  |  |  |  |
| NBS(-)LMP(-) |  |  |  |  | • | • | • | • | • |
| NBS(-)LMP(+) |  | • |  |  | • | • | • | • | • |
| NBS(+)LMP(-) |  |  |  | • | • | • | • | • | • |
| NBS(+)LMP(+) |  | • |  | • | • | • | • | • | • |

GA: gestational age; NBS: New Ballard Score; LMP: last menstrual period
\*Composite NBS: Sum of neuromuscular and physical maturity domains
<sup>^</sup>Birth weight 50%ile: Intergrowth 50th birthweight-for-age centiles used to convert birthweights to GA
<sup>†</sup>Individual NBS components: 5 Neuromuscular maturity domains and 7 physical maturity domains

The single parameter GA dating methods assessed include 1) LMP, 2) NBS, and 3) birth weight. GA dating by NBS was calculated from the composite NBS using the formula for GA conversion, as described by Ballard et al.[12] GA dating by birth weight was calculated under the naïve assumption that all infants are born at the 50^th^ birthweight‐for‐age centile and used INTERGROWTH standards[26] to convert these birthweights to GA.

The multiple parameter machine learning models assessed include 1) Optimized NBS, 2) NBS(‐)LMP(‐), 3) NBS(‐)LMP(+), 4) NBS(+)LMP(‐), and 5) NBS(+)LMP(+). In all machine learning models incorporating NBS, all 12 individual NBS components were included. With the exclusion of Optimized NBS, all machine learning models included five maternal and newborn parameters with various combinations of NBS and LMP. Maternal and newborn parameters included birth weight in addition to parameters with an *a priori* association with preterm birth (twin delivery, maternal HIV serostatus) and SGA (maternal height, maternal hypertension). We used hypertension in labor as a surrogate marker for maternal hypertension because, although imperfect, it is a readily accessible metric at delivery in the maternal delivery case file. Hypertension in labor was defined as systolic blood pressure ≥140 and/or diastolic blood pressure ≥90 recorded in the maternal delivery case file. HIV serostatus was determined by rapid ELISA performed according to local protocol at first antenatal care visit.[27]

We used super learner[28] to generate five GA and prematurity prediction models (components described above). In brief, super learner is a machine learning approach for combining the strengths of multiple predictive models or learners. Super learner finds the weighted, convex combination of these algorithms that minimizes the cross‐validated mean squared error of predictions of GA and preterm birth. To reduce concerns about over‐fitting the data, we utilized k‐fold cross validation (with 10 folds) to select the combination of learners. K‐fold cross validation ensures that the learner is not fit (trained) to the same data that are used to make predictions and judge performance. We used super learner computational macro (arXiv:1805.08058 [stat.ML]) developed in SAS version 9.4 (Cary, North Carolina) along with the SAS procedures HPFOREST, GENMOD, and GAM?. For our continuous GA prediction modeling of super learner models, we combined linear regression, random forest regression, and generalized additive models. For binary preterm birth classification modeling, we combined logistic regression, random forest classification, and generalized additive model.

Kernel density plots and Pearson’s correlation coefficients were generated to compare the predicted GAs from each continuous outcome model to GAs by early ultrasound. Receiver operating curves (ROCs) were generated and area under the curve (AUC) calculated for the diagnostic accuracy of preterm birth for each binary classification model. We also calculated the sensitivity, specificity, and percent correct classification for the identification of preterm infants using the best cutoff point for each model, as determined using the Youden method.[29] The Youden method determines a cutoff point by optimizing the differentiating ability of a test or model when equal weight is given to sensitivity and specificity. Subsequent sensitivity analyses were conducted on our best performing model to further assess stability and validity. All super learner modeling was performed in SAS as described above; all other analyses were performed using STATA release 14 (College Station, TX). This study was approved by the University of Zambia Biomedical Research Ethics Committee and the University of North Carolina Institutional Review Board.

## RESULTS

Between August 2015 and September 2017, 1450 pregnant women were consented and enrolled into the ZAPPS cohort. To date, 862 (59.4%) participants have had live births with deliveries captured by a study midwife. A total of 458 (53.1%) of these live births had newborns assessed at <72 hours of life by a trained nurse midwife and had complete data available to be included in subsequent preterm birth predictive modeling (Table 2). Among assessed live births, median ultrasound‐based GA was 39.6 weeks (IQR: 38.4‐ 40.3), with preterm birth prevalence 6.8%. The median birth weight was 3100g (IQR: 2855‐3400). The prevalence of SGA in this population was 14.1%. NBS assessment was the most common missing parameter, causing study exclusion (n=300; 76.1% of live births not assessed).

**Table 2:**
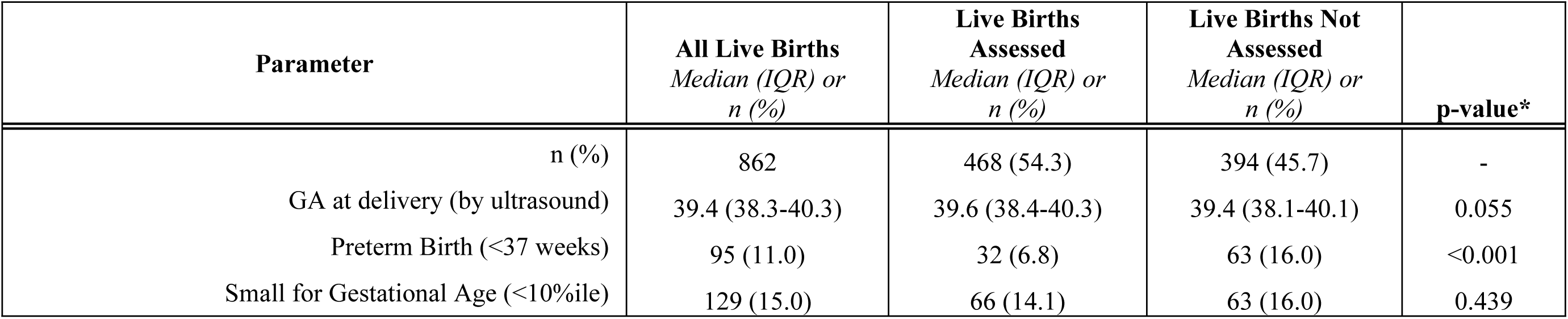
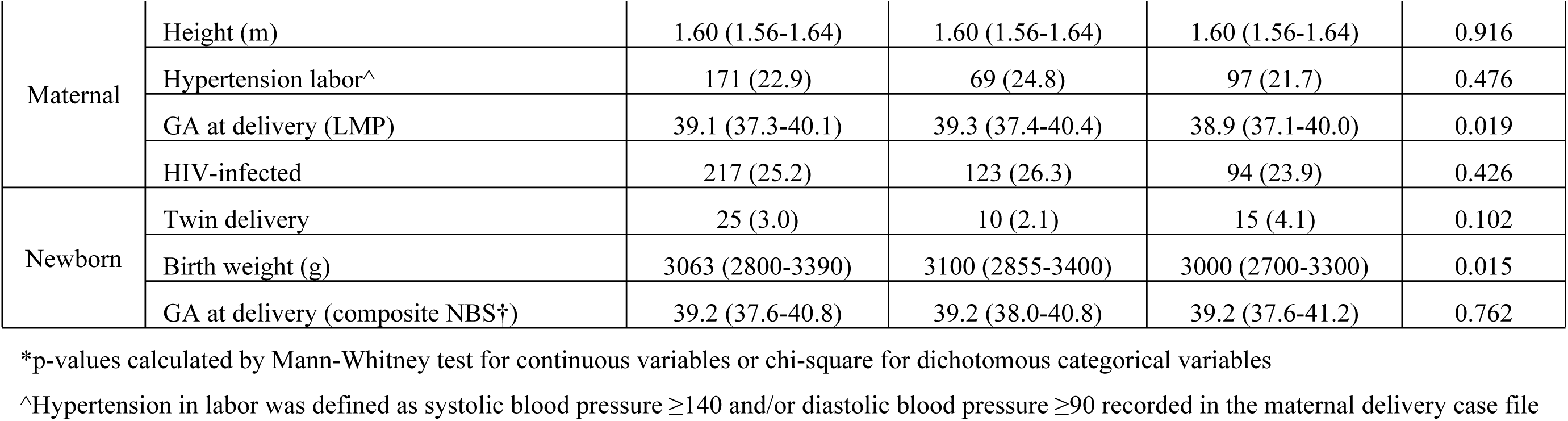
Maternal and newborn parameters included in gestational age modeling

| Parameter | All Live Births<br>Median (IQR) or<br>n (%) | Live Births<br>Assessed<br>Median (IQR) or<br>n (%) | Live Births Not<br>Assessed<br>Median (IQR) or<br>n (%) | p-value* |
| --- | --- | --- | --- | --- |
| n (%) | 862 | 468 (54.3) | 394 (45.7) | - |
| GA at delivery (by ultrasound) | 39.4 (38.3-40.3) | 39.6 (38.4-40.3) | 39.4 (38.1-40.1) | 0.055 |
| Preterm Birth (<37 weeks) | 95 (11.0) | 32 (6.8) | 63 (16.0) | <0.001 |
| Small for Gestational Age (<10%ile) | 129 (15.0) | 66 (14.1) | 63 (16.0) | 0.439 |

|  |  |  |  |  |  |
| --- | --- | --- | --- | --- | --- |
| Maternal | Height (m) | 1.60 (1.56-1.64) | 1.60 (1.56-1.64) | 1.60 (1.56-1.64) | 0.916 |
|  | Hypertension labor<sup>^</sup> | 171 (22.9) | 69 (24.8) | 97 (21.7) | 0.476 |
|  | GA at delivery (LMP) | 39.1 (37.3-40.1) | 39.3 (37.4-40.4) | 38.9 (37.1-40.0) | 0.019 |
|  | HIV-infected | 217 (25.2) | 123 (26.3) | 94 (23.9) | 0.426 |
| Newborn | Twin delivery | 25 (3.0) | 10 (2.1) | 15 (4.1) | 0.102 |
|  | Birth weight (g) | 3063 (2800-3390) | 3100 (2855-3400) | 3000 (2700-3300) | 0.015 |
|  | GA at delivery (composite NBS<sup>†</sup>) | 39.2 (37.6-40.8) | 39.2 (38.0-40.8) | 39.2 (37.6-41.2) | 0.762 |

\*p-values calculated by Mann-Whitney test for continuous variables or chi-square for dichotomous categorical variables
<sup>^</sup>Hypertension in labor was defined as systolic blood pressure $\geq 140$ and/or diastolic blood pressure $\geq 90$ recorded in the maternal delivery case file

Kernel density plots, with accompanying Pearson correlation coefficients, comparing the continuous GA distributions of the 8 models evaluated in this study to those calculated by ultrasound are shown in Figure 1. The models generated by the super learner program for GA as a continuous outcome clustered estimated GAs around the mean, resulting in a loss of outliers and less accurate estimation of GA as a continuous outcome (Figure 1d‐1h). Despite clustering around the mean, the NBS(‐)LMP(+) and NBS(+)LMP(+) machine learning models (Figure 1f, 1h) were found to best approximate the distribution of GA at delivery as compared to ultrasound dating (Pearson correlation coefficients 0.73 and 0.77, respectively).

**Figure 1:**
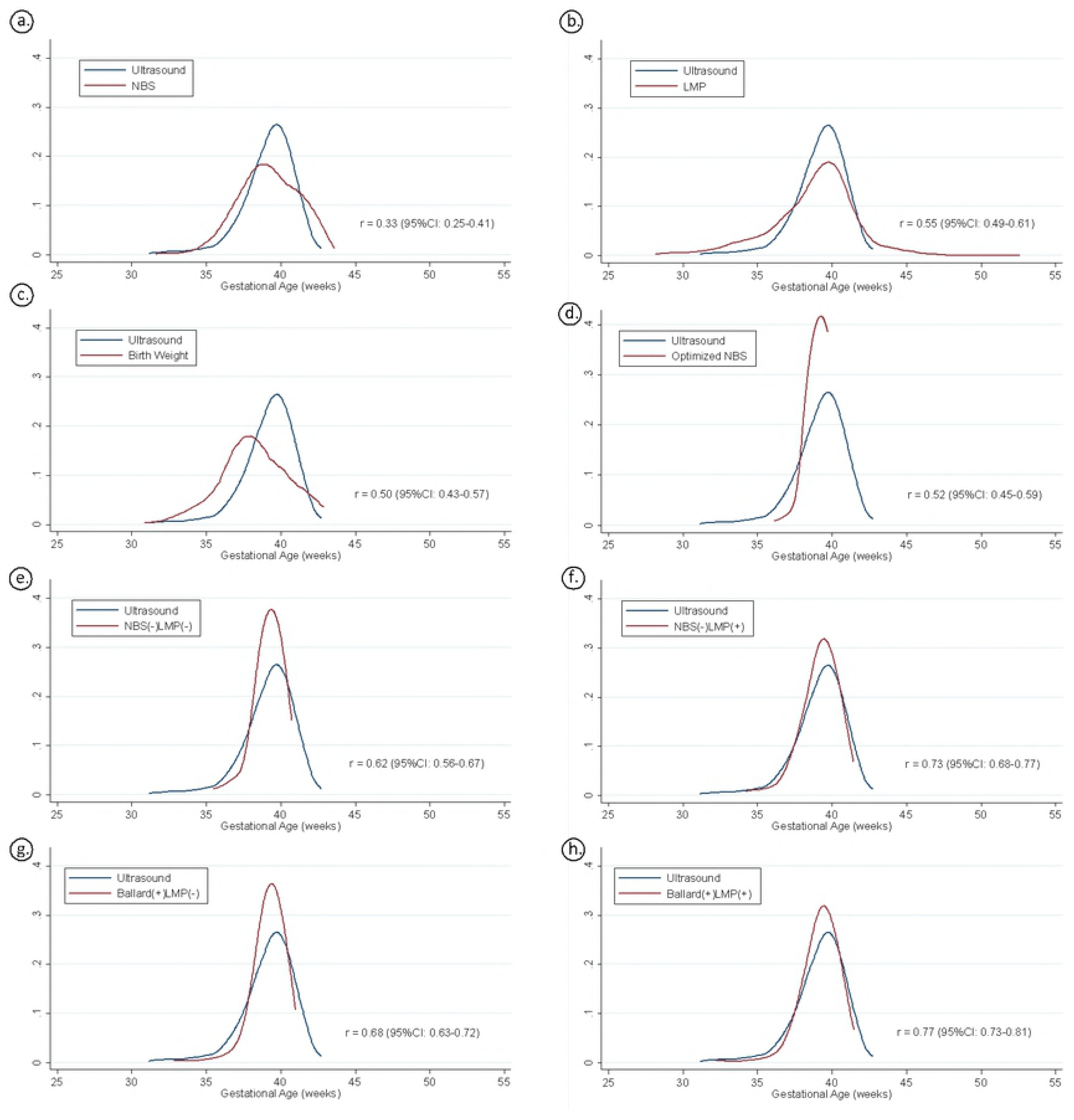
Distribution of gestational age at birth by all continuous models. r = Pearson’s correlation coefficient

The accuracy of preterm birth classification by each GA dating method was assessed using ROCs and associated AUCs (Figure 2). The AUC for Optimized NBS using super learner (0.8684) was improved compared to the single parameter NBS model (0.7645). The NBS(‐)LMP(‐) super learner model incorporating maternal and newborn parameters without LMP and NBS had an AUC of 0.8664, outperforming the NBS model and performing similarly to Optimized NBS. Adding LMP to this model, NBS(‐)LMP(+), improved the AUC (0.9796) more than adding NBS or both NBS and LMP (0.9242 and 0.9784, respectively).

**Figure 2:**
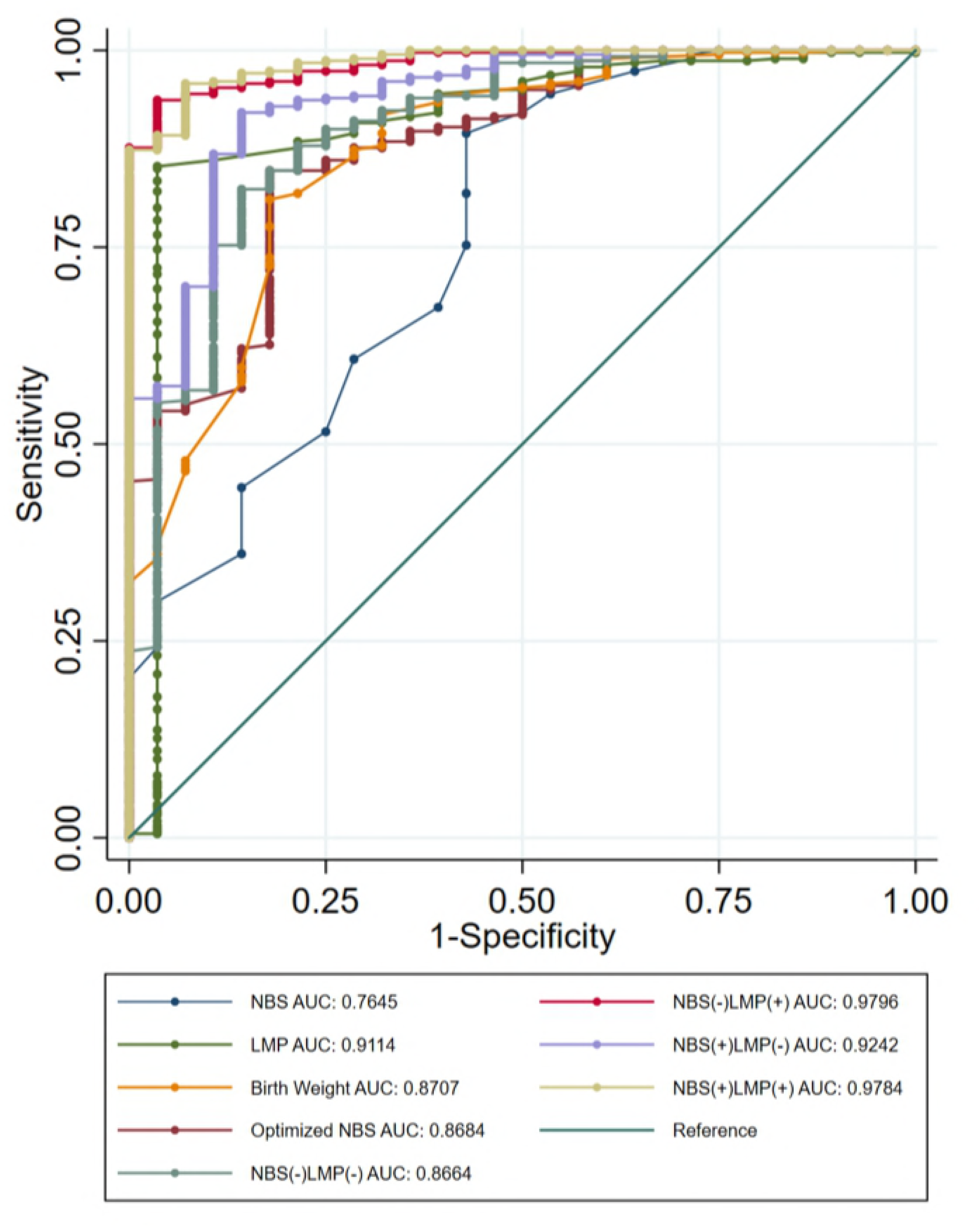
Diagnostic accuracy of binary models to identify preterm newborns. AUC: Area Under Curve

Sensitivity, specificity, and correct classification of all models predicting prematurity are shown in Table 3. In concordance with our ROC analysis, the NBS(‐)LMP(+) super learner model incorporating LMP without NBS had the highest percent correct classification (94.0%). The sensitivity and specificity achieved by this model were 93.8% and 94.0%, respectively.

**Table 3:** Prediction of preterm birth by binary models

| GA Dating Method | Sensitivity | Specificity | Correct Classification |
| --- | --- | --- | --- |
| NBS | 51.6% | 90.6% | 88.0% |
| LMP | 85.1% | 96.9% | 85.9% |
| Birth Weight | 82.6% | 75.0% | 82.1% |
| Optimized NBS | 81.3% | 85.3% | 85.0% |
| NBS(-)LMP(-) | 81.3% | 83.7% | 83.6% |
| NBS(-)LMP(+) | 93.8% | 94.0% | 94.0% |
| NBS(+)LMP(-) | 87.5% | 86.7% | 86.8% |
| NBS(+)LMP(+) | 96.9% | 88.1% | 88.7% |

Best cutoff point for calculations by each GA Dating Method determined using the Youden method

In sensitivity analyses, we tested NBS(‐)LMP(+), our best performing model, on the subset of women who were enrolled in the first trimester (<14 weeks; n = 204), as their ultrasound dating is expected to be most accurate, and we found a sensitivity of 96.6%, specificity of 94.3%, correct classification of 94.6%, and AUC of 0.9679. Additionally, because our best performing model excluded NBS – the variable most likely to cause study exclusion due to missingness – we were able to perform a second sensitivity analysis including 255 additional newborns on whom NBS was not collected. In this population of 713 children, the NBS(‐)LMP(+) model had a sensitivity of 91.3%, specificity of 86.7%, correct classification of 87.2%, and AUC of 0.9525.

## DISCUSSION

In our urban Zambian cohort with early pregnancy ultrasound dating, we used machine learning to identify a parsimonious set of maternal and newborn variables associated with prematurity and small for gestational age that can improve discrimination between preterm and term newborns as compared to common gestational age dating methods (New Ballard Score, last menstrual period, and birth weight). This exploratory study demonstrates the promising utility of machine learning techniques to optimize algorithms for the identification of preterm birth and other adverse birth outcomes in low‐resource settings.

Although a positive correlation between the number of parameters and accuracy of GA assessment has been established,[15] increasing parameter collection has negative feasibility of use, particularly in LMIC settings. In sub‐Saharan Africa, up to one‐half of all deliveries occur outside of the hospital and have no skilled birth attendant,[30],[31] limiting the utility of GA dating methods requiring numerous maternal and newborn metrics and characteristics. A significant strength of our best‐performing model is that it incorporates only six maternal and newborn characteristics and metrics available at delivery: LMP, birth weight, twin gestation, maternal HIV serostatus, hypertension at delivery, and maternal height.

An interesting finding of our analysis was the strength of LMP as a predictor of GA and preterm birth. Limitations of LMP as a dating method are well‐documented.[9] Women with lower educational attainment[9, 32] and later presentation to care[8, 33] tend to have less accurate recall of LMP, and the measure is subject to number preference (e.g. rounding to zero or five, or preference for 1^st^ or 10^th^ of month) and recall bias.[9, 10] Consequently, GA estimates by LMP alone suffer imprecision, with some estimates differing by weeks when compared to ultrasound.[34–37] Indeed, data from the Zambia Perinatal Record System, an electronic system that captured more than 250,000 births over a 6 year period in Lusaka, suggests an impossibly high preterm birth rate of 35% when LMP is used to determine GA.[11, 38, 39] Despite these limitations, we demonstrate that LMP is a useful predictor of prematurity and GA at delivery when incorporated into a model that allows obviously implausible estimates to be overridden by other parameters.

Further, our best performing prematurity prediction model excluded NBS. In fact, the addition of NBS components to our best performing list of covariates decreased the percent correct classification and AUC of the model. LMP outperformed NBS in prematurity prediction, both when assessed individually and in combination with other parameters. This finding supports a recent systematic review indicating that NBS has lower agreement with ultrasound dating than LMP.[15] The exclusion of NBS from our best‐performing model has the benefit of omitting lengthy and technical neonatal assessment procedures. Despite only including one newborn measurement, our model achieves an excellent AUC and correctly classifies more than 94% of newborns as preterm and term in our well dated Zambian cohort. Implementation of this model using six accessible maternal and newborn characteristics may increase the accuracy and rapidity of preterm newborn identification in LMIC settings as well as decrease the time and level of training required by frontline health workers to assess preterm birth.

A significant limitation of this current study is survival bias of assessed newborns, as demonstrated by the significant differences between our assessed and not assessed populations (Table 2). Preterm, especially early preterm, newborns were sometimes not assessed by study midwives because they were deemed too ill for the assessment or because of parental or neonatal provider objection to the exam. These early preterm newborns would likely have been identified as preterm by models included in this analysis. Thus, our estimates likely underestimate the performance of preterm birth identification in all models. Even with continuous staffing of the labor ward by midwives trained on NBS performance, many newborns were not evaluated within 72 hours. As many early preterm and critically ill newborns are never assessed, newborn assessments may not be the most effective measure of GA for these babies. In our cohort, we were able to assess significantly more newborns in our model (81% vs. 54%) when we included newborns on whom NBS was not collected, indicating that GA dating methods excluding newborn assessment may be more efficacious in LMIC settings.

A further limitation of our current model is that it was developed to optimize the accuracy of a binary outcome, preterm (<37 weeks) versus term (≥37 weeks) newborns, resulting in limited accuracy of our continuous GA dating models. Utilizing super learner capabilities to better model GA as a continuous outcome may be a helpful next step for neonatal providers desiring to better estimate accurate GA. Additionally, our current model requires all characteristics and measurements to assess preterm birth status be present for study inclusion. Consequently, if a woman does not know her LMP, her newborn is omitted from this model. Future work to assess novel GA and preterm newborn prediction models using machine learning techniques should include methods to impute missing data. Further, as validation of our model was limited to internal k‐fold cross validation, preventing over‐fitting the data, external validation should be pursued in future work.

In summary, by leveraging the capacity of cutting‐edge machine learning algorithms and maternal parameters associated with prematurity and SGA newborns, we identified a parsimonious list of covariates that improves accuracy of preterm newborn identification. Our model incorporates six accessible maternal and newborn characteristics and metrics, reducing the skill and time required to assess gestational age. This exploratory study supports the need for further research into the use of machine learning techniques to improve the accuracy of gestational age assessment in low resource settings and to assist frontline health workers in identifying newborns who may require special care.

